# Single-cell spatial multi-omics molecular pathology enabled by SuperFocus

**DOI:** 10.64898/2025.12.26.696575

**Authors:** Yao Lu, Xiaolong Tian, Marco Vicari, Archibald Enninful, Shuozhen Bao, Zhiliang Bai, Chen Liu, Xuchen Zhang, Per E. Andrén, Joakim Lundeberg, Mina L. Xu, Rong Fan, Yang Xiao, Zongming Ma

## Abstract

Histopathology and molecular pathology are currently distinct diagnostic modalities for the most part, one revealing tissue morphology at cellular resolution and the other providing molecular measurements with limited or no spatial context. Projecting genome-scale molecular information onto histopathology images at single-cell resolution across whole tissue sections represents a long-sought goal for next-generation pathology. Here we present SuperFocus, a modality-agnostic computational platform that generates histopathology-integrated single-cell spatial multi-omics from spot-based spatial measurements acquired on the same or an adjacent section without requiring external reference data. SuperFocus combines constrained cascading imputation with feature-level and cell-level quality-control scores to reduce spurious predictions and quantify confidence. On a ground-truth spatial transcriptomics benchmark dataset, SuperFocus improves key accuracy metrics by 28–73% over existing methods. Across Patho-DBiT, spatial ATAC–RNA, spatial CITE-seq and Visium-MALDI-MSI (SMA) datasets, SuperFocus enables cell-resolved analyses of MALT lymphoma microenvironments, gene regulatory programs in human hippocampus, lipotoxic hepatocyte states in human MASH, and transcriptomic-metabolomic states linked to neurotransmission and neuroinflammation in Parkinsonian mouse brain. Overall, SuperFocus enables scalable whole-slide single-cell spatial multi-omics integrated with histopathology, bridging histology and genome-scale molecular profiling for next-generation molecular pathology.

## Introduction

Spatially resolved molecular profiling has become indispensable for dissecting tissue organization, cellular interactions and disease heterogeneity *in situ* (1–7). By preserving anatomical context, spatial omics provides information that is fundamentally inaccessible to dissociated single-cell assays. Yet current technologies still impose practical trade-offs among experimental cost, spatial resolution, molecular breadth, and tissue coverage (8, 9). Spot-level measurement platforms such as Visium (10), DBiT-seq (11), spatial epigenomic (12), spatial CITE-seq (13), and MALDI-MSI (14) assays offer scalable and information-rich readouts, but their measurements are typically aggregated over spots or pixels that contain multiple cells. As a result, cell-level heterogeneity and whole-slide molecular architecture remain only partially resolved.

Histological images provide complementary information at cellular and tissue scales and remain the substrate of routine pathology. This makes the combination of spot-based molecular measurements and paired histology a particularly attractive foundation for computational refinement. Recent studies (15–17) have shown that tissue morphology can be integrated with spot-level molecular profiles to sharpen spatial maps and recover finer-scale molecular structure. However, the central challenge in molecular pathology is not simply to generate visually sharper maps. Existing morphology-guided refinement methods have been developed and benchmarked predominantly in transcriptomic settings, can fall short of robust cell-resolved inference, and generally lack built-in in-sample diagnostics to flag unreliable predictions. These limitations become more consequential when one seeks to generalize beyond transcriptomics to epigenomic, proteomic, metabolomic modalities and multi-omics settings, or to extend predictions beyond the profiled field of view across whole-slide tissue sections.

To address these challenges, we developed SuperFocus, a modality-agnostic computational platform that infers omics features for individual cells across whole-slide histology images from spot measurements acquired on the same or an adjacent tissue section, without requiring external reference data. SuperFocus is designed to operate across diverse spatial assays and to support cell-level spatial multi-omics analyses when modalities are co-profiled on matched or distinct spatial grids. In this way, SuperFocus is aimed not merely at super-resolution visualization, but at scalable molecular pathology over entire tissue sections.

At the method level, SuperFocus combines constrained cascading imputation across spatial scales with built-in quality control at both the feature and cell levels. The cascading design stabilizes prediction as resolution increases and enables molecular inference at the level of individual cells as well as intermediate subspot resolution levels. To quantify reliability, SuperFocus introduces feature-level and cell-level diagnostic scores that flag global and local deviations in prediction fidelity, as well as a per-cell credibility score that identifies extrapolative predictions outside the profiled region. Together, these components are designed to reduce spurious fine-scale structure, calibrate confidence and make cell-resolved whole-slide molecular inference more actionable in practice.

We systematically benchmarked SuperFocus on a high-resolution spatial transcriptomics dataset with single-cell ground truth. Across two feature-prediction accuracy metrics, SuperFocus improved performance by 28–73% relative to existing methods, while faithfully recapitulating cell-type composition and spatially localized biological programs. Importantly, the built-in quality-control measures tracked ground-truth-based validation closely and distinguished reliable from unreliable predictions with high sensitivity.

We then demonstrated the utility of SuperFocus across four molecular pathology settings spanning distinct tissues, disease contexts and assay modalities. In MALT lymphoma profiled by Patho-DBiT (18), SuperFocus resolved heterogeneous cell compositions and local cell–cell interaction programs across tumor microenvironments within a single tissue section. In a human hippocampus spatial epigenome–transcriptome dataset (19), SuperFocus inferred cell-level chromatin accessibility profiles that supported accurate cell-type annotation, spatial motif analysis, and the reconstruction of gene regulatory programs together with SuperFocus-resolved gene expression profiles on the same section. In human liver MASH profiled by spatial CITE-seq, SuperFocus-resolved cell-level protein maps identified a lipotoxic cell population that matched a metabolically stressed hepatocyte state defined independently from SuperFocus-resolved transcriptomic profiles. Finally, in a Parkinsonian mouse brain section profiled by Visium transcriptomics and MALDI-MSI on mismatched grids (20), SuperFocus enabled fine-grained cell-level integration of transcriptomic and metabolomic information across the tissue section, revealing cell populations and pathway signatures associated with dopamine depletion and spatially localized metabolic signals associated with local enrichment of microglia. Together, these results establish SuperFocus as a general, confidence-aware computational platform for whole-slide, cell-resolved molecular pathology from scalable spatial assays.

We have implemented SuperFocus as a Python package which is available at https://github.com/yaolu53/SuperFocus.

## Results

### SuperFocus enables whole-slide cell-resolved spatial multi-omics integrated with histopathology

We start with an overview of SuperFocus (**Super**-resolution re**f**inement of spatial **o**mics data with constrained **c**ascading imp**u**tation and whole-**s**lide cell-level prediction). The framework is modality agnostic. It can be deployed when the omics measurement is transcriptomics, epigenomics, proteomics, metabolomics, etc., as well as any combinations of them.

The input to SuperFocus includes two parts (**Fig. 1a**): (1) a high-resolution (20X or 40X) digitized histological (H&E) image of a tissue section, and (2) spot-level spatial omics measurements on the imaged section (or an adjacent one). Here, the omics measurements are possibly only within some relatively small but representative region(s) of the section. For example, the digitized H&E image could be obtained from whole slide imaging with 40X magnification and the spot-level omics data is obtained from applying some spatial omics technology on a selected region of interest (ROI) or multiple ROIs. When more than one sections are involved, we register all sections and impose a common coordinate system to align the histological image and the omics spots.

**Figure 1:**
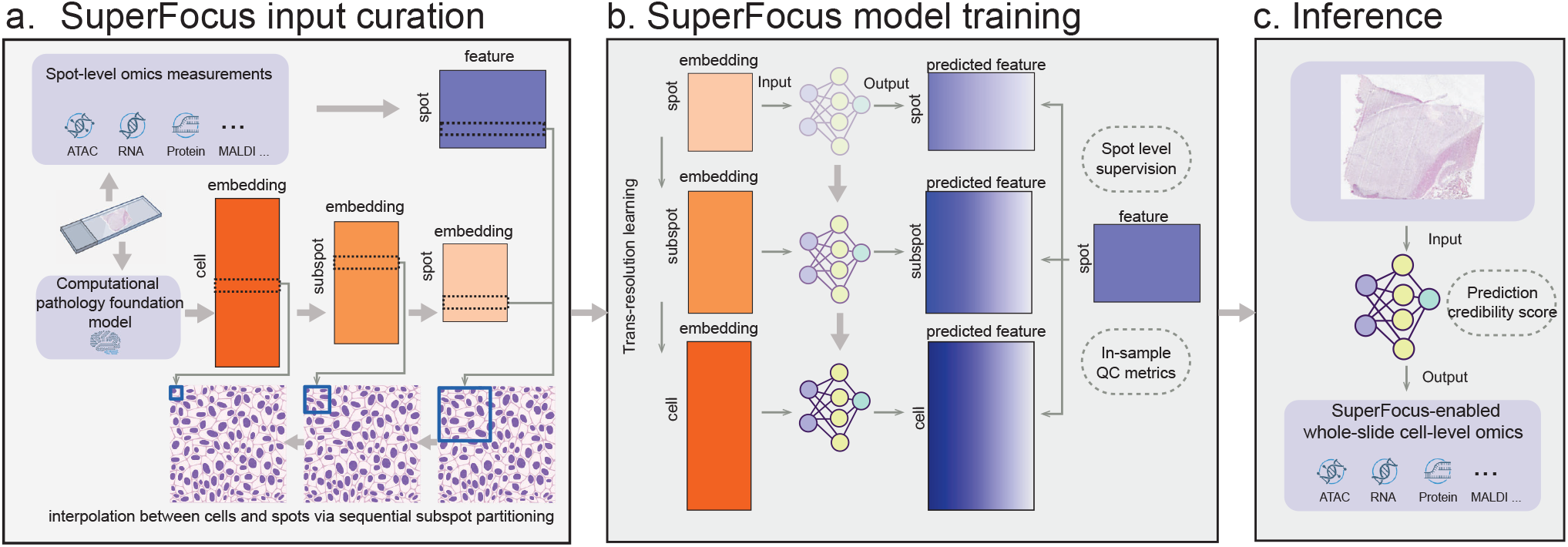
Overview of the SuperFocus pipeline. SuperFocus enables whole-slide cell-level inference from spot-based measurements across modalities by integrating histology. **(a)** In Stage 1 of SuperFocus, a computational pathology foundation model segments cells and encodes histology information for individual cells on the H&E image. SuperFocus curates spot-, sub-spot and cell-level features across different resolution levels that are suitable for predicting omics expressions. **(b)** In Stage 2, SuperFocus trains a cascade of prediction models with identical architecture at increasingly finer resolution levels via constrained cascading imputation, starting with the original spot level and ending with single-cell resolution level. In each round of training, the trained model parameter in the previous round serves as the warm start to enable trans-resolution learning. After training, in-sample quality-control metrics are computed to assess the performance of the trained SuperFocus model at predicting individual features. **(c)** In Stage 3, SuperFocus predicts cell-level expression profiles over the whole slide. Prediction-credibility scores for cells outside profiled FOV are computed to quantify cell-level prediction quality.

In Stage 1 of SuperFocus, we apply a computational pathology foundation model to segment cells and to encode cell morphological information provided by the histological image (**Fig. 1a**). By default, we use CellViT (21) in this work, while alternative models can also be used. We then reduce the dimensionality of the initial single-cell level CellViT image encoding by singular value decomposition. Next, we partition each omics measurement spot in a dyadic fashion recursively. While we use dyadic subspotting by default, other subspotting rates can also be used. To this end, we call the spatial resolution of the initial measurement spot the original resolution, and call the original spots resolution-level-0 spots. In each round of partitioning, we divide each spot at the current resolution level into four disjoint subspots of finer spatial resolution, and we call these subspots resolution-level-*ℓ* subspots where *ℓ* is the counter of dyadic partitioning round. We stop the recursive process when the majority of subspots contain only one cell. Hence, the finest resolution level is effectively at single-cell level. After recursive partitioning, we compute image encoding vectors for each (sub)spot at each resolution level thus created by aggregating the encoding of individuals cells within the (sub)spot and the immediately surrounding (sub)spots at the same resolution level. See Materials & Methods for additional details. Moreover, we count the number of cells within each (sub)spot at each resolution level. For omics data, we apply the standard pre-processing suitable for each modality.

In Stage 2 of SuperFocus, we define a cascade of predictive models, one for each resolution level (**Fig. 1b**). At each resolution level, the goal of the corresponding predictive model is to predict the average single-cell level omics expression of each (sub)spot at the level using the (sub)spot’s image encoding vector. The deep neural network architectures of these predictive models are intentionally designed to be identical in order to facilitate transfer learning (22) across resolution levels. Starting at level 0 (i.e., the original omics spot resolution), we train the cascade of predictive models recursively, with learned parameters of the previous resolution level model serving as a warm start of the training process of the current model to achieve trans-resolution learning. For the precise definition of these models and the tuning parameters used in training them, see Materials & Methods.

To evaluate the capacity of the trained model for predicting individual features, we conclude Stage 2 by computing two in-sample diagnostic (quality control) metrics for each omics feature. First, in each original omics spot, we sum up the predicted feature values by the trained finest resolution model across all finest resolution subspots and compute the difference of the sum from the observed feature value at this spot. We call this difference the aggregated prediction error of the spot and we collect all the aggregated errors across original omics spots into a long vector whose length equals the number of original spots. The first metric, relative sum-of-squared-errors (RSSE), is the ratio of the sum of squared aggregated errors over the sum of squared observed values across original spots, which aims to reflect the global in-sample prediction errors made by the finest resolution model. Higher RSSE values mean worse overall predictions. The second metric, error spatial locality (ESL), is the ratio of the *ℓ*_2_ norm of the aggregated error vector over its *ℓ*_1_ norm, which measures how concentrated the aggregated error vector is. Higher ESL values mean more spatially concentrated prediction error patterns. If the trained model at the finest resolution level (i.e., the trained model at the effective single-cell resolution level) predicts a feature well, both metrics should be low for that feature. As both metrics are unit-free, they can also be used to compare prediction accuracy across different features. We note that both metrics are in-sample, i.e., their evaluations only require the training data, and hence they can always be computed when SuperFocus is applied. Moreover, both metrics generalize directly to all resolution level at which the SuperFocus model has been trained.

Stage 3 of SuperFocus leverages the trained model at the finest resolution (i.e., the effective single-cell resolution level) to predict omics features for all individual cells over the entire imaged tissue section (**Fig. 1c**). To this end, for each cell, we first create a 3-by-3 array of subspots of the same size as the finest resolution ones, with the central subspot centered at the cell center. Next, we compute the image encoding of the central subspot which aggregates all image encoding vectors of all cells intersecting with the 3-by-3 array of subspots. This serves as the input to the trained finest-resolution model. Finally, we use the resulting output of the trained finest-resolution model as the predicted omics features of the cell. We repeat the foregoing process for all cells to generate single-cell omics features over the whole slide. For each cell predicted, we further assign it a prediction-credibility score that decreases with the distance in morphological space from its local neighborhood to those of training data. A low prediction-credibility score for a cell suggests that the histological patterns surrounding the cell is undersampled or absent from the training data, and hence one should not be overly confident in its predicted omics profile. The SuperFocus-predicted omics profiles for individual cells across the whole slide are not merely higher-resolution visualizations. They provide a quality-controlled, analysis-ready, cell-resolved molecular representation of tissue architecture and support a wide range of downstream analyses, as we exemplify in the rest of this section. Since SuperFocus maps molecular information onto a common histology-defined cellular canvas, it also creates a general interface for cross-resolution and cross-modality integration of spatial omics data acquired on the same or adjacent sections, as illustrated by the transcriptomic–metabolomic analysis in **Fig. 6**. More broadly, it can bridge spot-based spatial assays to external single-cell references in other modalities, thereby enabling downstream diagonal integration with multimodal methods such as MaxFuse (23) and GLUE (24). These analyses underscore that SuperFocus functions not only as a refinement method, but also as a general platform for quality-controlled, cell-resolved spatial multi-omics analysis for molecular pathology.

### Benchmarking and reliability validation of SuperFocus on a ground-truth VisiumHD human kidney dataset

We first benchmarked SuperFocus on a human kidney VisiumHD data (25) with single-cell level ground truth, which enabled validation with the same measurement technology on the same tissue section (**Fig. 2a**). The VisiumHD data was binned to 64 *μ*m spots (i.e., spots of size 64 × 64 *μ*m^2^) for curating spot-level training dataset, comparable to (albeit slightly larger than) the typical spot sizes in Visium and DBiT-seq experiments. On the other hand, we binned the VisiumHD data at 8 *μ*m spots for curating the validation dataset. Thus, the validation set was effectively of single-cell resolution and directly binning at 8 *μ*m resolution avoided potential confounding induced by cell segmentation in the validation set. Overall, we had 10,504 non-empty 64 *μ*m spots in the training set and 128,002 non-empty 8 *μ*m spots in the validation set. After preprocessing and screening based on the training data, a panel of 2,010 spatially highly variable and kidney marker genes were retained in both training and validation. Both SuperFocus and all methods in comparison used the same training set and were validated on the same validation set.

**Figure 2:**
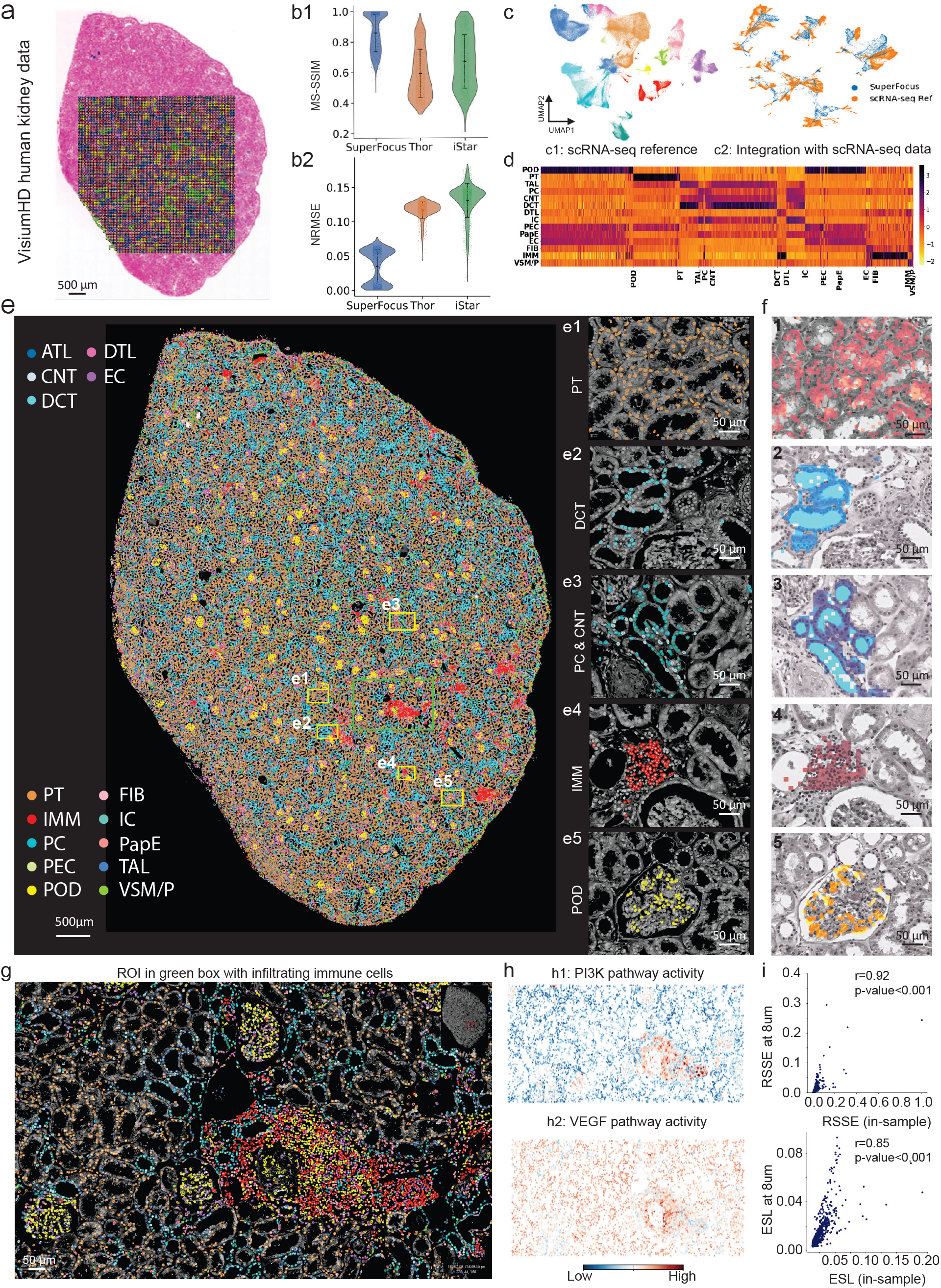
Benchmarking SuperFocus on a VisiumHD human kidney dataset. **(a)** High-resolution H&E image of human kidney sample with VisiumHD experiment region, spot size: 8 *μ*m. **(b)** Top: Side-by-side violin plots of MS-SSIM scores of SuperFocus (left: mean 0.86, SD 0.08), Thor (middle: mean 0.59, SD 0.16), iStar (right: mean 0.67, SD 0.18) predicted expression levels and ground-truth VisiumHD measurements across 2,010 spatially highly variable and kidney cell marker genes. The mean MS-SSIM score of SuperFocus improves that of Thor by ∼46% and that of iStar by ∼28%. Bottom: Side-by-side violin plots of NRMSE scores of SuperFocus (left: mean: 0.035, SD 0.024), Thor (middle: mean 0.117, SD 0.012), iStar (right: mean 0.131 SD 0.025) predicted expression levels and ground-truth VisiumHD measurements across the same 2,010 genes. The mean NRMSE score of SuperFocus improves that of Thor by ∼70% and that of iStar by ∼73%. **(c)** UMAP visualizations of scRNA-seq reference data from healthy human kidney samples (left) and of Harmony-integrated scRNA-seq reference data and SuperFocus-predicted cell-level gene expression profiles (right). **(d)** The heatmap of average SuperFocus-predicted gene expression profiles shows distinct patterns across cell types. **(e)** Gloabl and local spatial plots of annotated cell types of SuperFocus-predicted cell expression profiles. E1: proximal tubular cells, E2: Distal convoluted tubular cells, E3: Collecting duct cells, E4: Immune cells, E5: Podocytes. **(f)** F1 to F5 are corresponding regions to E1 to E5. The highlighted regions are distinct clusters from VisiumHD clustering results based on 8 *μ*m data by 10x Genomics. **(g)** A zoomed-in view of the region highlighted in green in panel **E. (h)** Spatial plots of two pathway activity scores: PI3K (top) and VEGF (bottom). **(i)** Plots of in-sample diagnostic metrics RSSE (top) and ELS (bottom) computed from 64 *μ*m spot training data show high correlations with ground-truth error metrics of overall prediction error and error spatial locality at 8 *μ*m spot resolution level.

We trained SuperFocus along a sequence of 64, 32, 16 and 8 *μ*m spots. We stopped at 8 *μ*m as 80.62% of the 8 *μ*m subspots contained no cell and 92.57% of the non-empty 8 *μ*m subspots overlapped with just one cell. The predicted gene expressions at each resolution level by the trained SuperFocus model at the respective resolution level were compared to the binned VisiumHD data at the same resolution level to inspect the multi-resolutional performance of SuperFocus. We found that SuperFocus accurately predicted gene expressions across different resolutions (**Supplementary Fig.1& 2**). For instance, the transcript *NPHS2*, which provides instructions for making podocin and expressed exclusively in podocytes, solely enriched within the glomeruli of the kidney. In addition, the transcripts *SLC34A1* and *SLC4A1*, which are canonical markers for cells in the proximal tubule and distal parts, respectively, showed congruent spatial patterns between SuperFocus predictions and VisiumHD bins across resolutions. We first benchmarked gene expression prediction accuracy of SuperFocus against iStar (16) and Thor (17), two state-of-the-art spatial resolution enhancement methods for spatial transcriptomics, on the 8 *μ*m resolution validation dataset using two quantitative accuracy metrics. The first metric, multi-scale structural similarity score (MS-SSIM) (26) with base pixel size at 8 *μ*m, quantifies the spatial similarity between SuperFocus prediction and ground truth for each gene. For each gene, a higher MS-SSIM score indicates better prediction, with 1 being the highest possible score. The second metric, normalized root-mean-squared-error (NRMSE) (17), provides a unit-less measure of the overall deviation of predicted values from the ground truth. For each gene, a lower NRMSE score indicates better prediction, with 0 being the best possible score. See Materials & Methods for their precise definitions. The side-by-side violin plots in **Fig. 2b** demonstrated the sizable improvements achieved by Super-Focus over existing methods in both metrics. The mean MS-SSIM score over 2,010 genes of SuperFocus (0.86) improved that of Thor (0.59) by ∼45% and that of iStar (0.67) by ∼28%. The mean NRMSE of SuperFocus (0.035) improved that of Thor (0.117) by ∼70% and that of iStar (0.131) by ∼73%.

Next, we benchmarked SuperFocus in its capacity of cell state differentiation with its predicted cell-level transcriptomic profiles. To this end, the trained SuperFocus model at the finest resolution level was applied to the H&E image of the whole tissue section to conduct *in silico* prediction of gene expression profiles for each of the 130,134 cells identified on the entire H&E image, including both cells within the profile ROI and those outside. To annotate the SuperFocus-predicted cell-level transcriptomic profiles, we integrated them with a reference single-cell RNA-seq dataset (27) using Harmony (28). The reference dataset consisted of 107,701 cells across 15 major cell types from 31 healthy donors (**Fig. 2c1**). After Harmony integration, SuperFocus-predicted expression profiles formed well-separated clusters which in turn aligned with major cell type clusters in the reference data (**Fig. 2c2**). The imperfect mixing was expected as scRNA-seq and VisiumHD are intrinsically different technologies. We annotated each SuperFocus-predicted profile using majority voting by cell types of its *k*-nearest neighbors in the reference dataset after integration. After cell type annotation, the heatmap of the average SuperFocus-predicted expression levels for marker genes of major kidney cell types showed distinct patterns (**Fig. 2d**). These patterns aligned well with known biology, with canonical marker genes predicted to have high expressions in the corresponding cell types, e.g., *PODXL* in podocytes. These findings validated the capacity of SuperFocus-predicted cell-level profiles in delineating cell types. Note that the foregoing integration with single-cell reference was not strictly necessary, as we could alternatively apply any chosen single-cell annotation method on the SuperFocus-predicted expression profiles directly.

Taking the validation of cell state delineation capacity of SuperFocus further, we benchmarked the SuperFocus annotations against the 8 *μ*m pixel clustering result published by 10x Genomics together with this dataset (25). To this end, we first mapped SuperFocus induced cell type annotations onto the tissue section, and found that the spatial distribution of the cell types accurately reflected the kidney anatomical structure (**Fig. 2e**). For example, the apical surface of proximal tubule was covered with dense, long microvilli, forming a prominent brush border (**Fig. 2e1**), while the distal convoluted tubule (DCT) and the collecting duct system (PC, IC and CNT) had a smooth apical membrane (**Fig. 2e2 & e3**). In addition, immune cells resided between the tubules (**Fig. 2e4**), and podocytes were located within glomerulus (**Fig. 2e5**). When compared with the ground-truth 8 *μ*m spot cluster labels, these distinct spatial niches with cell-level resolution enabled by SuperFocus refinement of 64 *μ*m spot-level data aligned nearly perfectly with distinct ground-truth 10x 8 *μ*m spot clusters (**Fig. 2f1-5**).

To further validate that SuperFocus enables cell-resolved downstream analyses, we zoomed in a region enriched with immune cells (**Fig. 2e** ROI within green bounding box & **Fig. 2g**). SuperFocus prediction showed that the enlarged glomerulus in this region, unlike other glomeruli, demonstrated inflammation and infiltration by immune cells, such as CD4+ T cells. This was supported by the higher activity score of the PI3K signaling pathway within this glomerulus (**Fig. 2h1**), which is a major regulator of cellular processes in podocytes, particularly when under stress and inflammation (29). In addition, we observed elevated activities through the hypoxia, the JAK-STAT and the MAPK signaling pathways (**Supplementary Fig. 3**). The glomerulus contains the most active and crucial VEGF signaling axis in human kidney (**Fig. 2h2**), which is fundamental to maintaining the glomerular filtration barrier. These SuperFocus-prediction-based spatial pathway activity plots exhibited high concordance with their ground-truth-based counterparts, along with multiple other examples (**Supplementary Fig. 3**).

Finally, we validated the informativeness of the feature-level in-sample quality-control metrics, RSSE and ESL, using this high-resolution VisiumHD dataset. Using only the 64 *μ*m spot training data, RSSE and ESL were strongly associated with their ground-truth counterparts computed by comparing SuperFocus predictions to VisiumHD measurements across 8 *μ*m spots (**Fig. 2i**; RSSE (top): *r* = 0.92, P-value *<* 0.001; ESL (bottom): *r* = 0.85, P-value *<* 0.001). To compute the ground-truth metrics for each gene, we constructed a vector of true prediction errors across all 8 *μ*m spots. The ground-truth RSSE is the ratio of the sum-of-squared values of this error vector over the sum-of-squared VisiumHD observed values over all 8 *μ*m spots. The ground-truth ESL is the ratio of the ℓ_2_ norm of this vector of true 8 *μ*m spot prediction errors over its *ℓ*_1_ norm. See Materials & Methods for details. In sum, with the high-resolution VisiumHD data, we systematically benchmarked and validated SuperFocus in its cell-level prediction accuracy (with ∼ 28% to ∼ 73% improvements over state-of-the-art methods in key metrics), in its cell state delineation accuracy, in its capacity in enabling down-stream analyses, and in the informativeness of the built-in quality-control metrics.

See **Supplementary Fig. 4** for additional validation results on a paired human breast cancer Visium–Xenium dataset on adjacent sections with single-cell level ground-truth. In particular, the validation of the effectiveness of the built-in prediction-credibility score for informing out-of-FOV cell-level prediction quality.

### SuperFocus unveils intratumoral cell-cell interaction programs in MALT lymphoma

We used SuperFocus to predict single-cell RNA profiles on a MALT lymphoma (mucosa-associated lymphoid tissue lymphoma) within a gastric mucosa H&E tissue section (30). Patho-DBiT measurements were obtained on a nearby tissue section (**Fig. 3a1**). The two sections were aligned by STalign and with the square in **Fig. 3a1** indicating the aligned Patho-DBiT FOV. The SuperFocus models were trained with the H&E image and 5,293 20 *μ*m Patho-DBiT spots (5,293 out of the original 10,000 spots contained at least one cell identified by Cell-ViT), with the finest resolution level set at 10 *μ*m. At 10 *μ*m, 46.38% subspots contained no cell and 77.99% non-empty subspots intersected with only one cell. The trained SuperFocus model at the finest resolution level was subsequently applied on the H&E image to predict gene expression profiles for a total of 157,689 cells identified by CellViT over the entire section.

**Figure 3:**
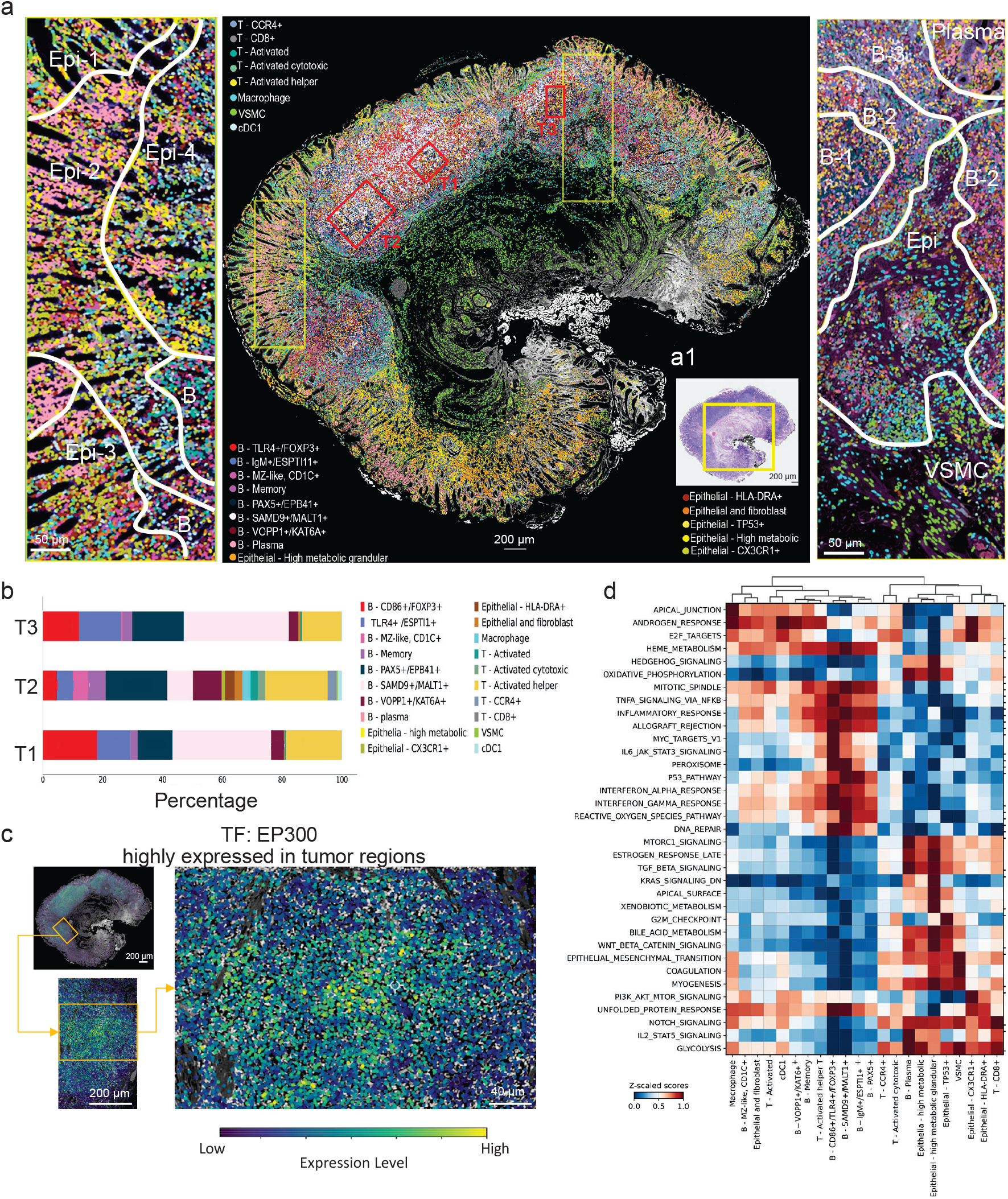
SuperFocus unveils intratumoral cell-cell interactions in MALT lymphoma with Patho-DBiT data. **(a)** Middle: Spatial plot of cells annotated by SuperFocus-predicted cell-level gene expression profiles; Left: A zoomed-in view of the ROI in the yellow box on the left peripheral region with epithelial cells as the dominating cell type; Right: A zoomed-in view of the ROI in the yellow box on the right with plasma cells, tumor B cells, VSMC, and epithelial cells as dominating cell types. **(a1)** High-resolution H&E image of MALT lymphoma sample with Patho-DBiT experiment region, spot size: 20 *μ*m. **(b)** Decomposition of SuperFocus-inferred cell types within tumor niches T1, T2 and T3. **(c)** Spatial plot of transcription factor *EP300* in a MALT lymphoma region (top left) with zoomed-in views at two levels (bottom left and right) **(d)** Hallmark gene set analysis results based on SuperFocus-predicted single-cell gene expression profiles.

SuperFocus-predicted RNA expressions aligned well with known pathological characteristics of the MALT lymphoma both within and outside of the region corresponding to the Patho-DBiT experiment FOV (**Supplementary Fig. 5**). For example, the transcript *MS4A1*, a marker for B cells, was enriched in the tumor regions, and the T cell marker *CD3D* concentrated around the periphery of follicles. The transcripts *XBP1* and *JCHAIN*, both of which are markers for plasma cells, were elevated in the lamina propria, where *MS4A1* expression level was low.

Next, we clustered the cells by applying a standard scRNA-seq data clustering pipeline on the SuperFocus-predicted expression profiles. Cell type annotation was then performed by taking into account both canonical gene markers and high-abundance transcripts within each cluster. See the middle panel in **Fig. 3a** for the spatial plot of the annotated cell types from the SuperFocus-predicted expression profiles and the two side panels for zoomed-in views of the two regions with yellow bounding boxes. Comparing the SuperFocus-enabled cell type annotations with the tissue architecture inferred from iStar (16, 30) (segmented by solid lines in the two side panels of **Fig. 3a**), we found that SuperFocus enabled the spatial study of this sample at a much finer granularity, allowing truly single-cell resolution delineation of different spatial niches and tumor microenvironments. For instance, in the left panel of **Fig. 3a**, within epithelial niches previously discovered (30), a composition of different epithelial cell subgroups were identified by SuperFocus, such as *TP53*+, *CX3CR1*+ and ones with high metabolic activities. In addition, memory B cells were present in the Epi-3 region, aligning well with the fact that this region was close to the tumor area. For the B cell niches, MZ-like and memory B cells were found across these peripheral follicle regions. Along with B cell subgroups, a group of activated T cells were also identified in this region. These findings agreed with the known biology about the organization of cell populations in lymph nodes. The majority of cells were plasma cells in both the region above Epi-3 on the left panel and plasma region on the right panel, consistent with previous findings and with the spatial plot of *JCHAIN* expression levels in **Supplementary Fig. 5**. The B-2 and B-3 regions aligned well with the mantle zone of two adjacent B cell follicles, where a large group of memory, *IgM*+ and MZ-like B cells were present. Apart from the B-2 and B-3 regions, B cells in the B-1 region demonstrated up-regulation of genes that promote proliferation, such as *PAXA5* and *TLR4*, which is a characteristic of germinal centers.

We then inspected three tumor niches identified in (30) indicated by red bounding boxes in the middle panel of **Fig. 3a**. Tumor region T1, previously annotated altogether as *INPP5D*+ B cells by iStar (30), consisted of 5 major B cell subgroups and activated helper T cells (**Fig. 3b**). Moreover, a larger proportion of *FOXP3*+ B cells were identified by SuperFocus in this region. The *FOXP3* gene has been found to suppress immune response (31) and this finding was in accordance with the local high expression levels of the *INPP5D* gene, which acts as a negative regulator of B-cell receptor signaling in immune cells (32). Tumor region T3, characterized by enhanced B cells proliferation in (30), exhibited an increased proportion of *TLR4*+ B cells compared to other regions of interest (ROIs) (**Fig. 3b**). The *TLR4* gene is known to promote B cell proliferation and survival via the ERK, the AKT and the NF-*κ*B pathways (33). B cells with upregulated *EPB41* gene were found present by SuperFocus in this tumor niche (**Fig. 3b**). The *EPB41* gene plays a structural and regulatory role in connecting the membrane to the actin cytoskeleton in B cells (34), which aligned with the previous findings that B cells in this ROI underwent actin cytoskeleton organization (30). Notably, SuperFocus further delineated the complex cell type compositions within these exemplifying ROIs and identified the presence of activated helper T cells in all of them. Previous studies discovered, by using customized immunochemistry panels and high-resolution imaging, that the helper T cell population was more prevalent and highly active in MALT lymphoma compared to controls (35), and that these helper T cells play a promoting role in MALT lymphoma by creating a supportive microenvironment that enables malignant B cells to better survive and proliferate (36). As we have demonstrated, SuperFocus gained comparable insights at single-cell spatial resolution from spot-level data without a targeted panel, providing single-cell level details of iStar-annotated spatial niches beyond a single enriched cell type per niche.

Furthermore, we analyzed cell-cell communication patterns across different cell types in this MALT lymphoma sample from a hallmark pathway perspective using the SuperFocus-predicted single-cell gene expression profiles (**Fig. 3d**). Based on cell type annotations with SuperFocus-predicted transcriptomic profiles, MZ-like and memory B cells, macrophages, and cDC1 cells located in the periphery of follicles, other B cell subgroups except plasma cells along with activated helper T cells mainly resided in the central tumor regions, and the lamina propria consisted of different types of epithelial cells, plasma cells, smooth muscle cells, and some T cells. The Hallmark pathway analysis revealed that cells in the periphery of follicles, the central tumoral regions, and the lamina propria interacted via distinct pathways (**Fig. 3d**). For example, the NF-*κ*B and the JAK-STAT3 pathways, two of the most critical pathways in MALT lymphoma, were active in cells, especially *TLR4*+ B cells, located in the central tumor regions (37). Meanwhile, malignant B cells showed enhanced IFN-*α* and IFN-*γ* pathway activities, which are critical in the pathogenesis and progression of MALT lymphoma, particularly those associated with chronic infection with helicobacter pylori (38). Additionally, the mitotic spindle assembly, which directly links to the uncontrolled proliferation of malignant B cells, was found to be active in tumor regions. Furthermore, some pathways exhibited inconsistent activity scores in different tumor regions within this sample (**Supplementary Fig. 6**). These results not only illustrated a high level of heterogeneity within this tumor sample but also demonstrated the ability of SuperFocus to faithfully capture important biological signals in its prediction.

Finally, TF scoring results obtained from SuperFocus prediction identified TFs that were known to be more active in MALT tumors (**Supplementary Fig. 7**). In particular, this sample showed upregulated *EP300* expression (**Fig. 3c, Supplementary Fig. 7**). The EP300 transcription factor is a key player in the pathogenesis of B-cell lymphomas through its function as a major histone acetyltransferase (HAT) and transcriptional co-activator (39). Morevoer, *NOTCH1*, one of the known aberrant activators often linked to the pathogenesis of B-cell malignancies (40), was also predicted to have increased expression levels within the tumor regions (**Supplementary Fig. 7**). Additionally, SuperFocus enabled the examination of TF expression levels within each cell, identifying *TLR4*+, *IgM*+ and *ESPTI1*+ B cells and activated T helper cells in the central tumor regions with aberrant increase in both TFs, and warranted further investigation into how activated TFs mediated cells within tumor regions (**Supplementary Fig. 7**).

### SuperFocus infers single-cell chromatin accessibility from spot-based spatial epigenomic data and enables cellular gene regulatory network analysis

We evaluated SuperFocus on a human hippocampus co-mapping data, where high quality H&E image (**Fig. 4a1**) and spot-level spatial ATAC-seq and mRNA-seq co-profiling measurements (**Fig. 4a2**) were obtained from two adjacent sections (19). The tissue sections were aligned by STalign (41) with anchor points. We trained SuperFocus models using the H&E image and spot-level ATAC-seq data from 5,000 spots at the initial resolution 50 *μ*m first, and then at 25 *μ*m and 12.5 *μ*m. At the resolution 12.5 *μ*m, 85.09% subspots contained no cell, and 81.07% non-empty subspots overlapped with only one cell. After preprocessing, 32,802 peaks were retained from the spot-level data and used for SuperFocus training. The trained SuperFocus model at the finest resolution level was subsequently applied on the adjacent H&E image to predict the peak profiles for a total of 49,372 CellViT-identified cells.

**Figure 4:**
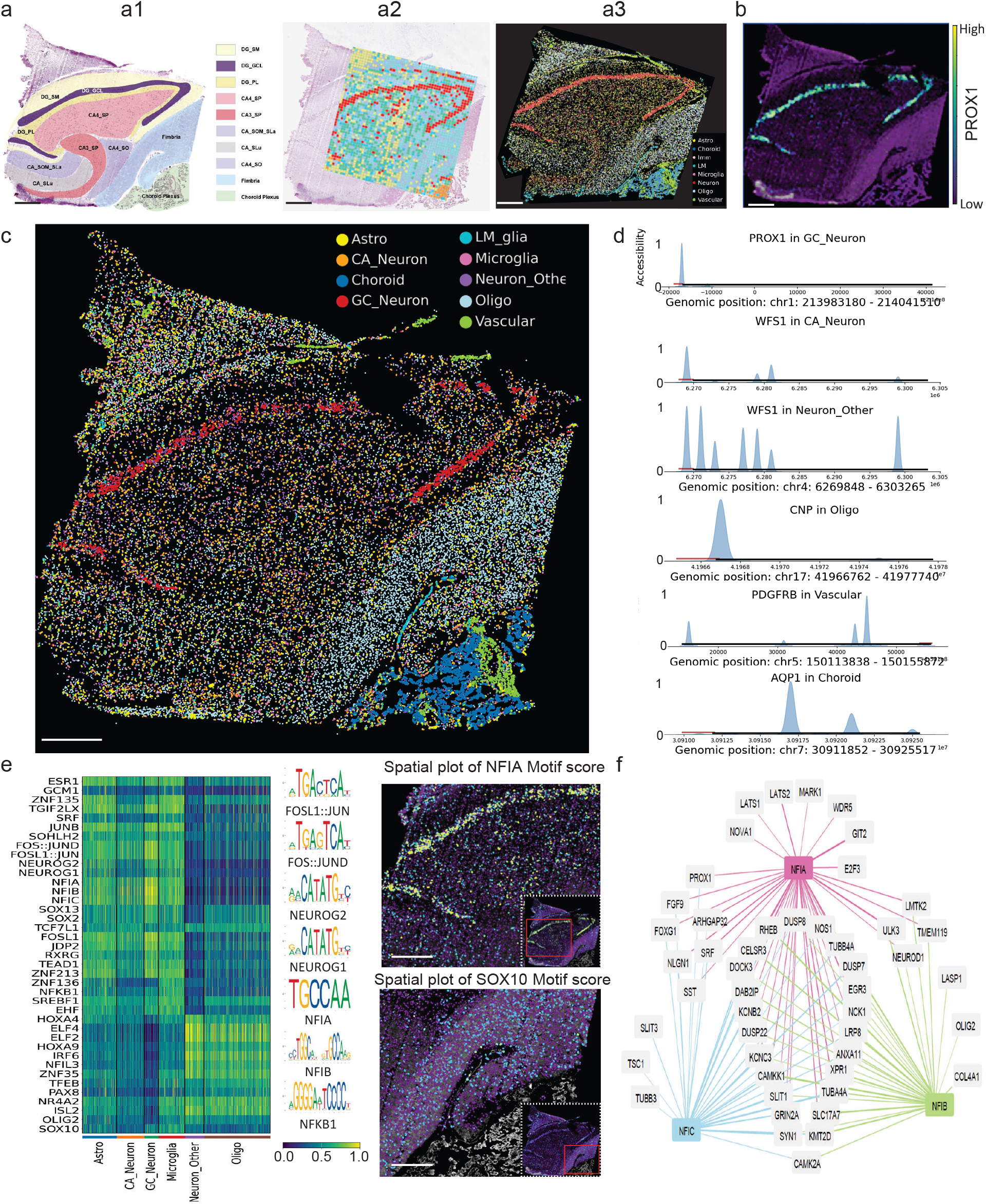
SuperFocus infers single-cell chromatin accessibility from spot-level human hippocampus spatial epigenomic data and enables cell-resolved spatial regulatory analysis via same-slide transcriptomic integration. **(a)** a1: High-resolution H&E image of the human hippocampus sample (scanned at 40× magnification by a slide scanner Leica SCN400, 0.25 *μ*m resolution); key histological components and layers were annotated by a neuropathologist. DG_SM: stratum moleculare of dentate gyrus; DG_GCL: granule cell layer of dentate gyrus; DG_PL: polymorphic layer of dentate gyrus; CA4_SP: stratum pyramidale in cornu ammonis 4; CA3_SP: stratum pyramidale layer in cornu ammonis 3; CA_SOM_SLa: somatostatin and stratum lacunosum in cornu ammonis; CA_Slu: stratum lucidum layer in cornu ammonis; CA4_SO: stratum oriens layer in cornu ammonis 4; Fimbria: fimbria-fornix complex. a2: Spot-level spatial ATAC-RNA-seq data for human hippocampus sample (only ATAC results shown here), spot size: 50 *μ*m; a3: Spatial plot of cell-type annotation, generated from 25-plex immunohistochemistry proteomic staining on an adjacent section. **(b)** Spatial plot of SuperFocus-predicted single-cell level *PROX1* gene expression. **(c)** Cell type annotation based on SuperFocus-predicted single-cell level peak profiles. **(d)** Pseudo-coverage plots of the gene regions for marker genes *PROX1, WFS1, CNP, PDGFRB, AQP1* in their corresponding cell types inferred from SuperFocus-predicted cell-level chromatin accessibility profiles. **(e)** Heatmap of TF deviation scores inferred from SuperFocus-predicted cell-level peak profiles across different cell types (left). Spatial plot of *NFIA* motif binding site accessibility across cells, focusing on neurons subgroups (top right). Spatial plot of *SOX10* motif binding site accessibility in oligodendrocyte region (bottom right). **(f)** Inferred gene regulatory network related to three spatially variable TFs, NFIA, NFIB and NFIC, based on paired SuperFocus-predicted cell-level gene expression and peak profiles. Scale bars: 1 mm.

Next, we conducted Leiden clustering of the SuperFocus-predicted single-cell peak profiles using standard single-cell ATAC-seq data protocol and annotated each cluster based on the peak-inferred gene activity scores (42) of marker genes (**Fig. 4c**). For instance, open chromatin profiles at the coding regions of *OLIG2 and CNP* gene identified oligodendrocytes, while *WFS1* and *PROX1* identified neurons. Immunohistochemistry (IHC) for key cell type–specific marker proteins was performed on adjacent sections by spatial proteomics (Akoya PhenoCycler, 25-plex) and used as ground truth for the spatial distribution of different cell types relative to the anatomy of human hippocampus and surrounding regions. A board-certified neuropathologist confirmed the identification of major cell types based on IHC results, including neurons, oligodendrocytes, astrocytes, microglia, vascular cells, leptomeningeal cells, and choroid plexus cells, with tissue contexts (**Fig. 4a3**).

Comparing **Fig. 4a3** and **Fig. 4c**, we found that SuperFocus predictions and IHC cell type annotations were highly concordant, and both aligned well with the anatomical organization of the human hippocampus (**Fig. 4a1**). SuperFocus-predicted peak profiles enabled us to accurately locate neurons in Dentate Gyrus (DG Neuron) and Cornu Ammonis CA Neuron), as well as oligodendrocytes (Olig) and astrocytes (Astro). Dentate gyrus neurons were densely packed within the granule cell layer, whereas astrocytes and oligodendrocytes were distributed across the molecular and polymorphic layers of the Dentate Gyrus. The fimbria were enriched for oligodendrocytes and largely devoid of neurons. Moreover, SuperFocus-predicted peak profiles successfully discriminated Choroid Plexus (ChPx), Vascular (Vas) and Leptomeningeal (LM) cells.

To further validate the biological faithfulness of SuperFocus-predicted single-cell epigenomic profiles, we examined the spatial distributions of SuperFocus-predicted mRNA expression levels of key marker genes by training a separate SuperFocus model on the mRNA-seq assay of this co-profiled spatial dataset. After spot-level spatial transcriptomic data pre-processing, a total of 2,026 transcripts were used in SuperFocus training and prediction. We found that the *PROX1* gene, a marker for mature granule neurons, was highly expressed in Dentate Gyrus granule cell layer (**Fig. 4b**). Moreover, *MOG* was enriched in oligodendrocytes, *SST* was expressed where neurons were identified, regions with high expressions of *SLC1A2* agreed with the locations of astrocytes, and *PECAM1* showed elevated expression level in vascular cells (**Supplementary Fig. 8**). The pseudo-coverage plot of SuperFocus-predicted peaks showed that the genome was more accessible within the *PROX1* gene region for Dentate Gyrus granule neurons than other cell types (**Fig. 4d, Supplementary Fig. 8**), that the *WFS1* gene region was more accessible in different neuron subgroups, and that accessibility peaks were found in regions related to marker genes of oligodendrocytes, vascular cells and choroid plexus (**Fig. 4d**).

Furthermore, the SuperFocus-predicted single-cell open chromatin peak profiles offers a new perspective to investigate TF binding activity across individual cells with spatial contexts in an intact tissue. We calculated the TF deviation score (43) to quantify accessibility at genomic regions containing specific TF binding motifs, relative to background levels. Highly variable TF motifs were then identified for each cell type, and a heatmap of standardized TF deviation scores was generated (**Fig. 4e**, left). Neurons in the Dentate Gyrus and Cornu Ammonis showed sizably higher chromatin accessibility in the genomic regions containing the motifs for the *FOSL1::JUN, FOS::JUN, NEUROG1* and *NEUROG2* transcription factors. Homeostatic microglia, activated microglia, and other resident immune cells of the central nervous system exhibited increased chromatin accessibility around the *NFKB1* TF motif region, consistent with elevated regulatory activity associated with the NF-*κ*B pathway. Alongside the heatmap, SuperFocus provided us the opportunity to view the motif binding activities spatially. *NFIA* motif binding activities restricted to various neurons, while the *SOX10* motif binding activities were mainly among oligodendrocytes (**Fig. 4e**, right).

SuperFocus enables the construction of cell-level gene regulatory networks from this spot-level co-profiling dataset (**Fig. 4f**). It integrated chromatin peak–gene associations, transcription factor activity, and motif occurrences to derive a quantitative score that quantifies the regulatory strength between transcription factors and their target genes. See **Materials & Methods** for details. For instance, SuperFocus constructed gene regulatory networks for the NFI family transcription factors (*NFIA, NFIB*, and *NFIC*), which are primarily associated with glial fate and the regulation of neuronal function (**Fig. 4f**) (44). *NFIA* and *NFIB* motif accessibility is high in astrocytes, microglia, and granule neurons (**Fig. 4e**). This pattern has been consistently reported in bulk and single-cell ATAC-seq studies of the human brain (45) and is concordant with the established roles of NFI transcription factors in gliogenesis and neuronal migration. *NFIC* has been shown to regulate distinct downstream neuronal genes, including *SST*, which modulates neuronal excitability, and *SLIT3*, which is involved in axon guidance and neurovascular development (**Fig. 4f**).

### SuperFocus resolves cell-level protein expressions *in situ* and identifies lipotoxic hepatocyte subpopulation validated by same-slide transcriptomics

We applied SuperFocus on a spatial CITE-seq dataset measuring human metabolic dysfunction-associated steatohepatitis (MASH) liver sample at 50 *μ*m (**Fig. 5a & b**). We first focused on the protein modality for hypothesis generation, with validation evidence obtained independently from the transcriptomics modality. To this end, we independently trained separate SuperFocus models for the proteomics and the transcriptomics assays.

**Figure 5:**
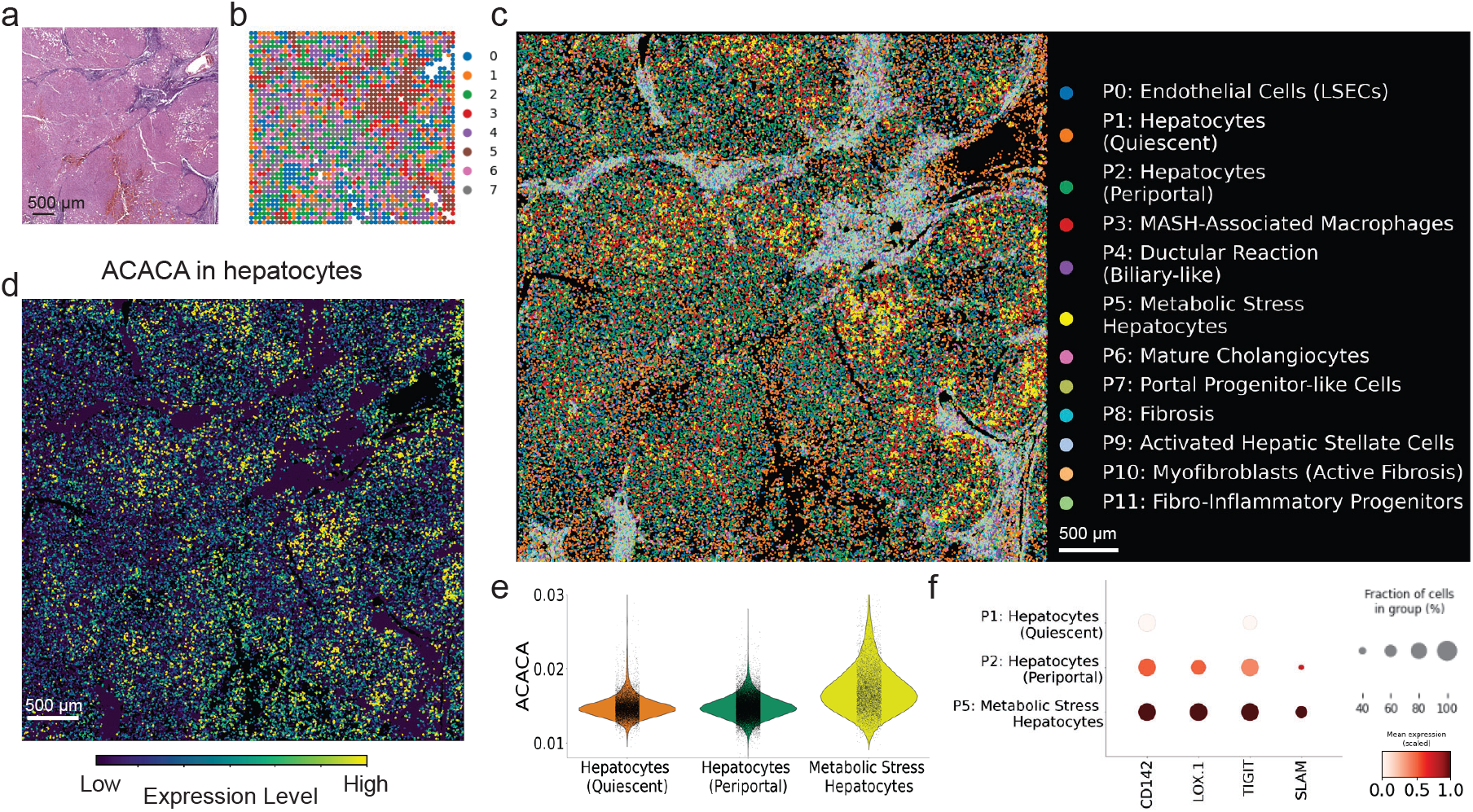
SuperFocus enables the identification and validation of a lipotoxic hepatocyte subpopulation in a human liver MASH spatial CITE-seq dataset. **(a)** High-resolution H&E image of the human MASH liver sample (scanned at 40× magnification, 0.25 *μ*m resolution); **(b)** Spot-level spatial CITE-seq data for MASH liver sample (only protein results shown here), spot size: 50 *μ*m; **(c)** Spatial plot of Leiden cluster labels (P0–P11) inferred from SuperFocus-predicted single-cell level protein profiles. Each cluster was then annotated by analyzing the highly expressed genes using single-cell level transcriptomic data predicted by an independently trained SuperFocus model on the RNA assay. **(d)** Spatial plot of the lipid-metabolism related gene *ACACA* in hepatocytes from SuperFocus-refined single-cell level transcriptomic data. **(e)** Violin plot of SuperFocus-refined *ACACA* expression level in three hepatocyte subgroups. **(f)** Dot plot of four stress-related protein markers from SuperFocus-refined protein data across three hepatocyte subgroups.

The protein assay consists of 163 markers in 2,483 spots, with 2,098 passing quality control in pre-processing and being used to train the protein SuperFocus models for this section. See **Fig. 5b** for spot-level cluster annotations based on observed spot protein expressions. The SuperFocus model cascade was trained till the spatial resolution reached 12.5 *μ*m. At 12.5 *μ*m resolution, 51.52% subspots contained no cell, and 74.30% non-empty subspots overlapped with just one cell. The finest-resolution model was then used to predict protein expressions for all 105,281 cells within the ROI. See **Fig. 5c** for the spatial distributions of cell-level clusters based on the predicted cell-level protein profiles. The clusters P0–P11 in **Fig. 5c** were defined solely by the SuperFocus-resolved cell-level protein profiles, and the displayed biologically meaningful cell type annotations after the colon marks for these protein clusters were obtained from further examining the SuperFocus-predicted transcriptomic profiles of the cells within each cluster later in the validation stage. While we included them directly in **Fig. 5c** for conciseness, the hypothesis generation stage used only the protein modality and we refer to the clusters only by the labels within this stage. Cells in the P5 cluster (colored in bright yellow) in **Fig. 5c** demonstrated an elevated level of the LOX-1 protein, a primary scavenger receptor for oxidized LDL (**Fig. 5f**). A high expression level of LOX-1 indicates a high level of oxidative stress and lipid-induced injury (46). In addition, upregulation of CD142 and TIGIT is a commonly observed response to inflammatory metabolic stress in MASH (47). A high expression level of SLAM, a protein involved in the crosstalk between hepatocytes and immune cells, is a sign of progression of chronic liver injury and fibrosis (48). All these proteomic signals suggested that this cell subpopulation suffered from lipotoxicity.

To validate this hypothesis generated by SuperFocus-resolved cell-level protein data, we trained SuperFocus separately on the RNA assay till spatial resolution reached 12.5 *μ*m and predicted the expression levels of 3,005 highly-variable genes for all 105,281 cells in this ROI. Cell clusters based on SuperFocus-refined protein data were then further annotated based on highly expressed genes from SuperFocus-refined transcriptomic data (**Fig.5c**, cell type annotations after colon marks). See **Supplementary Fig. 9** for a dot plot of SuperFocus-resolved marker genes in these protein-defined clusters. Three clusters with high expression of *MALAT1* and *FNBP1* were annotated as hepatocytes. Cells in these clusters were distributed in nodules spatially. In addition, the *LPP* gene was a marker gene for fibrosis and clusters with highly expressed *LPP* were restricted in the dark purple regions in H&E (**Fig. 5a**). These findings aligned well with the structural features in the H&E image. In particular, cells in cluster P5 (colored in yellow) showed signs of notable metabolic dysfunction with upregulation of the genes *HPR, COTL1* and *FOXB1* (49), which supported the earlier protein-marker-based hypothesis. We thus annotated this groups of cells as Metabolic Stress Hepatocytes. To further validate our findings, we inspected the canonical gene associated with lipotoxicity of hepatocytes, *ACACA*. **Fig. 5d** plotted the spatial distribution of *ACACA* expression levels and **Fig. 5e** compared its expression levels across the three protein-based hepatocytes subpopulations via side-by-side violin plots. As *ACACA* catalyzes the carboxylation of acetyl-CoA to malonyl-CoA, its upregulation within the Metabolic Stress Hepatocyte subpopulation indicated surplus of lipid species (50, 51), resonating strongly with the proteomic evidence (**Fig. 5f**) of active inflammation and metabolic dysfunction in this cell group.

### SuperFocus enables cell-level integrative transcriptomic-metabolomic analysis in a mouse model of Parkinson’s disease

Integrating metabolomic data with other omics is technically challenging because metabolites behave very differently from genes or proteins. Metabolites (e.g., lipids, amino acids, nucleotides) span a wide range of molecular structures and physicochemical properties, and no single analysis platform can comprehensively capture their spatial heterogeneity. We showcased SuperFocus to reveal brain metabolites, especially neurotransmitters, using MALDI-MSI data (FMP-10 derivatized, 3,005 spots at 100 *μ*m spatial resolution) co-profiled with the entire transcriptome (10x Visium, 3,120 non-empty spots at 55 *μ*m spatial resolution) on a brain section from a 6-OHDA mouse model of Parkinson’s disease (20) . The striatum in one brain hemisphere was dopamine-depleted, whereas the contralateral hemisphere was neurotypical (**Fig. 6a**). Both the introduction of the new metabolomic modality and the mismatched spot sizes between the transcriptomic and metabolic modalities posed new challenges in the current setting (**Fig. 6b**).

**Figure 6:**
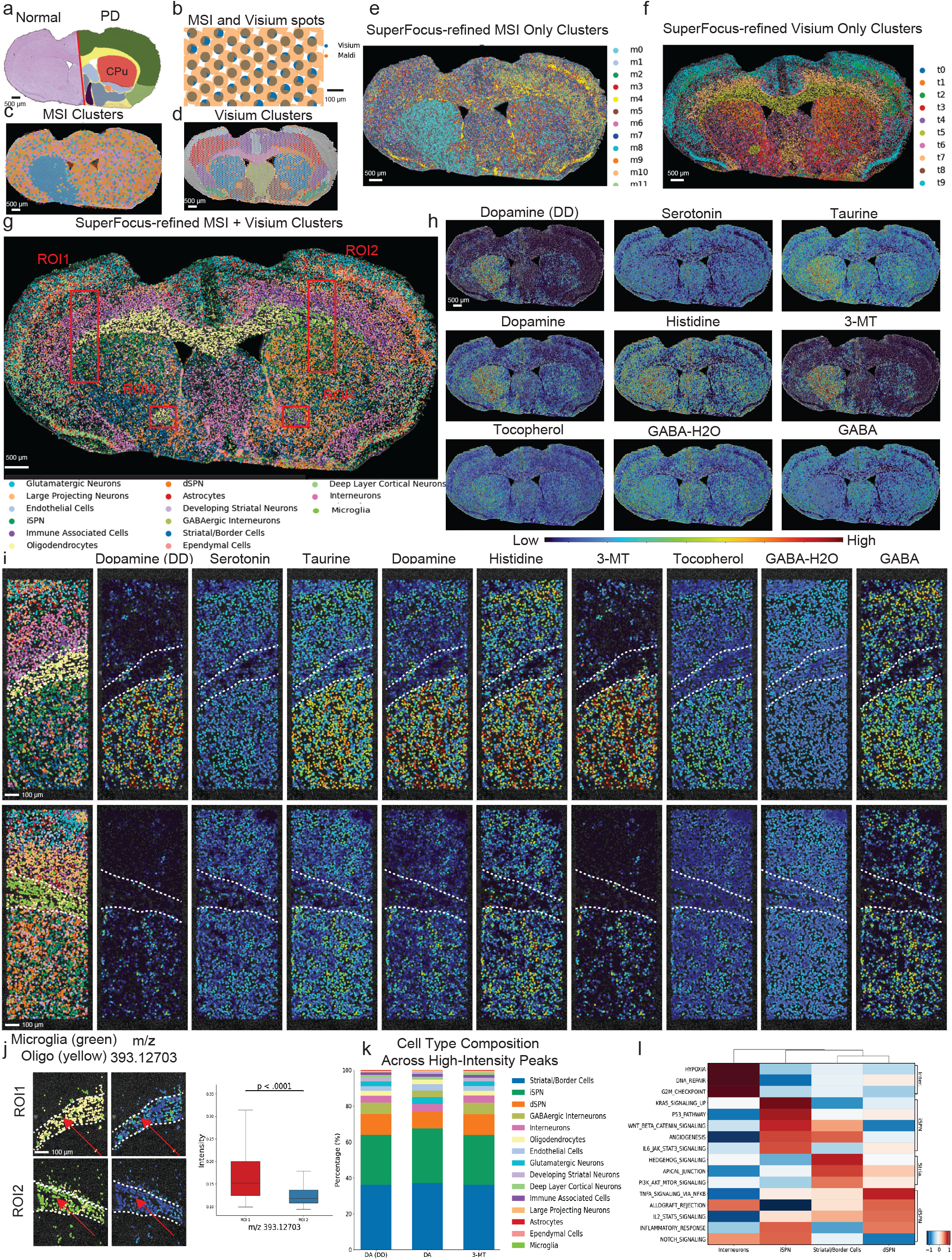
SuperFocus-enabled integrative transcriptomic-metabolomic analysis of a Parkinsonian mouse brain section. **(a)** High-resolution H&E image of the Parkinson diseased mouse brain (20x magnification, 0.5 *μ*m resolution). Key histological components and layers were annotated by a neuropathologist (dark green: cortex; light green: piriform cortex; dark purple: hopothelamus; dark blue: pallidum; light blue: striatum nucleus accumbens (bottom) and striatum leptal septal complex (left); orange: anterior commissure; red: caudate putamen (CPu); yellow: corpus callosum); **(b)** Overlay of the mismatching patterns of MALDI-MSI and Visium spots on the tissue, Visium spot size: 55 *μ*m, MALDI-MSI spot size: 100 *μ*m; **(c)** Clustering results based on 28 MALDI-MSI FMP-10 peaks at original MALDI-MSI spot resolution; **(d)** Clustering results based on 3,000 highly variable genes from Visium data at original Visium spot resolution; **(e)** Clustering results based on SuperFocus-refined 28-peak MALDI-MSI profiles; **(f)** Clustering results based on SuperFocus-predicted cell-level transcriptomic profiles; **(g)** Clustering results based on cell-level vertical integration of SuperFocus-refined MALDI-MSI and transcriptomic data by MOFA+, with cell type annotations based on the highly expressed genes in individual clusters (iSPN: indirect striatal projection neuron, dSPN: direct striatal projection neuron); **(h)** Spatial plots of neurotransmitters and metabolites from SuperFocus-refined MALDI-MSI data; **(i)** Zoomed-in views of cell type composition (left) and expression level of neurotransmitters and metabolites (right) in ROIs 1 and 2; **(j)** SuperFocus-annotated microglia (green) and oligodendrocytes (yellow) around the corpus callosum regions of ROIs 1 and 2 (left), SuperFocus-resolved signal intensities of taurine (m/z 393.12703) in the same regions (middle), side-by-side boxplots of m/z 393.12703 intensity level in cells in these regions (P-value *<* 0.001 in t-test for mean difference); **(k)** Compositions of cells with high expressions in DA(DD), DA, and 3-MT; **(l)** Pathway analysis in striatal border cells, dSPNs, iSPNs and interneurons.

We selected 28 m/z peaks within the MALDI-MSI data, which included neurotransmitters, such as dopamine, serotonin, and gamma-aminobutyric acid (GABA), and a group related to neuro-inflammation. See **Supplementary Fig. 10a** for details. The clustering result of the selected 28 m/z peaks at the original MALDI-MSI spot resolution was shown in **Fig. 6c**, which identified the dopamine-enriched area in the intact striatum in the normal hemisphere while missing other key structures of the mouse brain.

Leveraging the paired 20X H&E image, SuperFocus enhanced the spatial resolution of MALDI-MSI channels four times, from 100 *μ*m to 25 *μ*m in resolution. As suggested by the prediction accuracies on a hold-out validation set, 25 *μ*m spots offered a balance between resolution enhancement and prediction accuracy (**Supplementary Fig. 10b**). See **Materials & Methods** for details. The trained SuperFocus model at 25 *μ*m was then applied on the H&E image to predict average intensities of the peaks over a square spot of inscribed radius 12.5 *μ*m for all 64,680 cells on the tissue section. Leiden clustering of SuperFocus-refined m/z peaks identified regions in cortex, lateral septal complex and corpus callosum with different metabolism patterns while retaining the dopamine-enriched area (**Fig. 6e**). In addition, SuperFocus refinement enabled visualization of the MALDI-MSI peaks at enhanced spatial resolution (**Fig. 6h & Supplementary Fig. 11**), revealing spatial patterns consistent with the findings in (20) and comparable to those reported earlier on a 6-OHDA rat model with MALDI-MSI measurements (14).

To perform a cell-level integrative transcriptomic-metabolomic analysis, we further trained separate SuperFocus models on the Visium spot data from the same tissue section (starting at the original 55 *μ*m spots and with the finest resolution level set at 13.75 *μ*m) and thereby predicted the transcriptomic profiles of all 64,680 cells. At spatial resolution 13.75 *μ*m, 53.21% of the subspots were empty and 74.71% of the non-empty subspots intersected with just one cell. Compared with Leiden clusters of the original Visium spots, (**Fig. 6d**), Leiden clusters of SuperFocus-predicted data (**Fig. 6f**) distinguished distinct layers in the cortex region and different cell types in caudate and putamen (CPu). While neither the Leiden clusters of the original Visium spots nor those of the SuperFocus-predicted cell-level transcriptomic data distinguished the dopamine-enriched region, the SuperFocus-predicted cell-level transcriptomic data revealed notable shifts in the relative proportions of clusters t1 and t4 between the normal and lesioned hemispheres (**Fig. 6f**).

Having obtained cell-resolved transcriptomic and neurotransmitter profiles from SuperFocus, we next performed cell-level vertical integration of the two modalities using MOFA+ (52). Leiden clustering of the resulting joint embeddings, followed by marker-based annotation, recovered anatomically coherent cell populations across the section (**Fig. 6g**). For example, striatal border cells were concentrated near the dopamine-enriched boundary of the left CPu, whereas striatal projection neurons localized mainly within the CPu and formed two subpopulations corresponding to the direct and indirect pathways. The corpus callosum, by contrast, was enriched for oligodendrocytes, consistent with its white-matter identity. To examine lesion-associated changes, we defined two approximately symmetric ROIs spanning the cortex, corpus callosum, and CPu in the healthy and diseased hemispheres, respectively (ROIs 1 and 2 in **Fig. 6g & i**). Within these matched regions, the diseased-side CPu showed markedly reduced dopamine-related signals, whereas several metabolites, including serotonin, taurine, histidine, and GABA, were enriched in the cortex and CPu but sparse in the corpus callosum.

Through the integrative analysis, we further found that microglia were over-represented in the corpus callosum and anterior commissure on the lesioned side relative to the contralateral hemisphere (**Fig. 6j & Supplementary Fig. 10e**). This pattern is consistent with prior evidence that 6-OHDA lesions trigger microglial activation and neuroinflammatory responses in Parkinsonian models (53, 54), and with the emerging view that white-matter abnormalities contribute to Parkinson’s disease pathology (55). To investigate whether metabolomic changes in these regions were consistent with altered white-matter homeostasis, we examined the MALDI-MSI peak at m/z 393.12703, assigned to taurine in the FMP-10 dataset. Taurine is an aminosulfonic acid with neuromodulatory and homeostatic functions, and prior studies have linked taurine to oligodendrocyte precursor differentiation, myelin synthesis, and anti-inflammatory neuroprotection (56–58). The spatial plots and paired ROI comparisons showed significantly lower taurine intensity on the lesioned side in the corpus callosum (**Fig. 6j**), with a similar pattern in the anterior commissure (**Supplementary Fig. 10e**). Together, these findings are consistent with lesion-associated inflammatory and homeostatic remodeling in ipsilesional commissural white matter, potentially reflecting a reduction in a protective taurine-rich metabolic milieu in regions showing increased microglial representation (59, 60).

Finally, using the cell-level integrative profiles generated by SuperFocus, we identified cells associated with neurotransmitter signals enriched in dopamine-rich regions. In particular, we focused on three signals: Dopamine (double derivatized), Dopamine, and 3-methoxytyramine (3-MT). For each signal, we examined the cell-type composition of cells whose SuperFocus-resolved local average intensity exceeded the 95th percentile across all cells. We found that the majority of these cells were striatal/border cells, striatal projection neurons, and other interneurons (**Fig. 6k**). Notably, striatal/border cells, which accounted for roughly one-third of cells associated with elevated dopamine signaling, were spatially concentrated in dopamine-enriched regions and were under-represented in the lesioned hemisphere (**Supplementary Fig. 10c**). Further analysis of the hallmark pathways of striatal neurons and interneurons revealed that the Hedgehog pathway was particularly active in the striatal/border neurons (**Fig. 6l & Supplementary Fig. 10c**). Given prior evidence that Hedgehog signaling, particularly Sonic hedgehog derived from dopaminergic neurons, plays a critical role in maintaining striatal circuit integrity and modulating dopaminergic signaling balance (61, 62), these results were consistent with selective remodeling or loss of a striatal boundary neuron cell state in the lesioned hemisphere.

## Discussion

Spot-based spatial omics paired with histology provides one of the most practical routes to molecular profiling in intact tissue, as it offers rich molecular readouts at experimental scales that are difficult to achieve with uniformly cell-resolved assays and extends to multi-omics settings. The central limitation of this strategy, however, is that spot-level measurements blur cellular heterogeneity and usually cover only a restricted profiled region. In this work, we developed SuperFocus as a unified computational framework that converts these scalable inputs into whole-slide, cell-resolved molecular readouts. Rather than focusing only on visual super-resolution, SuperFocus is designed to recover molecular information for individual cells across tissue sections while retaining the capacity to explicitly quantify when predictions are likely to be unreliable. In this sense, SuperFocus shifts the practical operating point of spatial molecular pathology: it couples experimentally accessible spot-based assays with whole-slide, cell-resolved inference and built-in quality control.

The results that we have presented here support three aspects of this framework. First, benchmarking on a spatial transcriptomics dataset with matched single-cell ground truth showed that SuperFocus improves molecular feature prediction accuracy relative to existing methods, resulting in faithful recovery of cell states and spatially localized biological programs. Second, the built-in quality-control metrics tracked prediction fidelity closely as we have validated on the benchmarking study, indicating that reliability assessment can be incorporated as a core component of spatial molecular inference rather than treated as an afterthought. Third, the four application studies suggest that the SuperFocus paradigm is not limited to transcriptomic refinement, but can extend to epigenomic, proteomic, metabolomic, and multi-omic settings, including cross-modality integration on mismatched spatial grids. Together, these findings position SuperFocus not simply as another refinement method, but as a general computational route for extending scalable spot-based assays into cell-resolved molecular pathology.

Meanwhile, the present study also clarifies the conditions under which this route is likely to be most reliable. SuperFocus assumes that the profiled region captures histologic and cellular states that are sufficiently representative of the broader tissue section on which inference is performed. When the assayed region undersamples rare architectures, sharply localized programs or transitional states, prediction in unsampled regions becomes increasingly extrapolative. In this setting, the built-in quality-control scores are valuable as guardrails, but they do not eliminate the consequences of unrepresentative sampling. This makes ROI design an important upstream determinant of whole-slide inference quality. More broadly, it suggests that experimental design and computational inference should be treated as a coupled problem: the utility of whole-slide molecular reconstruction depends not only on model architecture and training strategy, but also on how well the profiled region spans the histologic and molecular diversities of the tissue.

This perspective creates a natural connection to recent advances in digital pathology foundation models and ROI selection methods. Large pathology foundation models can summarize tissue architecture across gigapixel slides, and emerging ROI-selection frameworks now use adjacent H&E sections and foundation-model features to prioritize regions that better capture histologic heterogeneity and molecular information content. Integrating these tools with SuperFocus could make whole-slide molecular reconstruction more systematic and more cost effective by guiding where spatial assays should be performed in the first place. In this view, SuperFocus is not only a downstream inference engine, but also part of a broader design loop in which histology guides assay placement and assayed regions support tissue-wide molecular reconstruction. Recent whole-slide pathology foundation models such as UNI (63) and Prov-GigaPath (64), together with ROI-selection frameworks such as S2-omics (65), make this combined strategy increasingly realistic.

Several additional limitations should be considered. The present benchmark is strongest in transcriptomic settings, so although the application studies support broader modality generality, each new assay class will still require modality-specific validation. This is beyond the scope of the current work, as public datasets that enable rigorous cell-level benchmarking outside transcriptomic spatial settings remain limited. Moreover, the quality-control scores introduced here should be interpreted as internal measures of reliability rather than substitutes for experimental truth. Prospective studies that combine SuperFocus with optimized ROI design and orthogonal validation assays will therefore be important for further assessing and validating its utility in translational and clinical molecular pathology workflows.

At a broader level, SuperFocus suggests that the longstanding trade-off between scalable acquisition and cell-resolved spatial analysis is not fixed at the level of experimental technology alone. By pairing standard spot-based assays with histology-guided inference with built-in quality control, it becomes possible to recover whole-slide molecular landscapes at cell-level granularity from experimentally accessible inputs. We anticipate that this framework will be useful not only for tissue-level biological discovery, but also for the emerging design of integrated molecular pathology workflows in which assay selection, ROI selection, computational reconstruction and downstream interpretation are optimized together.

## ACKNOWLEDGEMENTS

Computational data analysis was conducted with Yale High Performance Computing clusters (HPC). We acknowledge the support received from the U.S. National Institutes of Health including grants U54CA274509, U54CA268083, UH3CA257393, RF1MH128876, U54AG079759, U54AG076043, R01CA245313, RM1MH132648 (all to R.F.), and U01CA294514 (to R.F., M.X., and Z.M.), and from the U.S. National Science Foundation in grant DMS-2245575 (to Z.M.). We acknowledge support from the Single Cell Spatial Analysis Program (SCSAP) at the University of Michigan (to Y.X.). Cartoons in Figures 1 were created with BioRender.com. This article reflects the views of the authors and should not be construed as representing the views or policies of the institutions that provided funding.

## AUTHOR CONTRIBUTIONS

Conceptualization: Y.L., R.F., Z.M.

Algorithm Development and Implementation: Y.L., Z.M.

Analysis: Y.L., M.X., Y.X., R.F., Z.M.

Contribution of Material and Expertise: X.T., M.V., A.E., S.B., Z.B., C.L., X.Z., P.A., J.L., M.X., Y.X., R.F., Z.M.

Supervision: M.X, Y.X., R.F., Z.M.

